# Maternal separation remodels Purkinje cell circuits and impairs cerebellar synaptic plasticity

**DOI:** 10.64898/2026.11.08.765140

**Authors:** Pellegrino Lippiello, Eriola Hoxha, Claudia Cristiano, Silvia Pascarella, Floriana Volpicelli, Luisa Speranza, Mariella Cuomo, Maria Grazia Ferraro, Adrien Stanley, Anita Maria Rominto, Kardelen Dalim Filiz, Fabrizio Maria Liguori, Daniela Tropea, Silvia Di Angelantonio, Lorenzo Chiariotti, David Sulzer, Maria Concetta Miniaci

## Abstract

**Introduction:** Early-life adversity has been associated with cerebellar abnormalities, but the cellular and circuit-level changes involved and their persistence across development remain poorly understood. We examined whether maternal separation (MS) disrupts cerebellar circuit maturation and whether these changes are accompanied by alterations in synaptic function and behavior.

**Methods:** CD1 mice underwent daily maternal separation from postnatal day 2 to 18, whereas control litters remained undisturbed. Cerebellar alterations were assessed using transcriptomic and DNA methylation profiling, RT-qPCR, immunoblotting, Purkinje-cell morphometric analysis, and VGLUT1/VGLUT2 immunofluorescence. Patch-clamp electrophysiology was used to examine synaptic transmission and plasticity. Motor coordination, spontaneous locomotion, social approach, and stress-coping behavior were also evaluated.

**Results:** RNA sequencing of whole cerebellar tissue revealed early transcriptional remodeling, involving synaptic transmission, neuronal architecture, and plasticity-related pathways. Consistent with this profile, we found that MS altered Purkinje cell dendritic arborization and excitatory input remodeling, together with deficits in cerebellar long-term synaptic plasticity and reduced BDNF-related signaling. In parallel, methylation analysis revealed hypermethylation of the glucocorticoid receptor promoter and reduced cerebellar receptor expression, indicating that early-life stress also engages stress-related epigenetic mechanisms in this region. Importantly, these neurobiological changes were associated with motor coordination deficits, altered stress-coping behavior, and impaired social interaction.

**Discussion:** These findings identify the developing cerebellum as a target of early-life adversity and suggest that disrupted cerebellar maturation may contribute to lasting functional consequences. This study extends our understanding of the effects of maternal separation beyond the brain regions traditionally associated with stress responses.

## 1. INTRODUCTION

Maternal care plays an important role in the development of emotional and cognitive processes of offspring during the postnatal period. Studies in rodents have demonstrated that normal mother-pup interaction during the first two weeks of life ensures a reduced stress response in the pups, known as the stress hyporesponsive period (SHRP) (Levine, 1994; Weaver et al., 2004). This critical period is characterized by reduced corticosterone (CORT) levels as a result of enhanced glucocorticoid receptor-mediated negative feedback on the hypothalamic–pituitary–adrenal (HPA) axis. In humans, the SHRP is thought to begin around 6 months of age and to extend until 4–6 years of age (Gunnar and Donzella, 2002). Exposure to physical or psychological stressors during this sensitive period has been shown to impact adult behavior and increase vulnerability to neuropsychiatric disorders (Felitti *et al*, 1998; Nelson *et al*, 2020; Speranza *et al*, 2024).

In rodents, maternal separation (MS) is considered a reliable method for studying the physiological and behavioral effects of early life stress (Malave et al., 2022). MS induces hyperactivity of the HPA axis and increased release of CORT, thus exposing the brain to higher corticosterone levels (Rosenfeld et al., 1991). Glucocorticoids are known to affect neuronal structure, function, and synaptic remodeling, particularly during development, with long-lasting effects on brain plasticity and behavior later in life (Woolley et al., 1990; Liston and Gan, 2011). The early-life stress - induced increase in glucocorticoids may thus affect the maturation and functions of brain regions, such as the hippocampus, amygdala, and prefrontal cortex, implicated in stress adaptation and emotional regulation.

The cerebellum is a brain region that is potentially highly sensitive to the effects of early-life stress, as it exhibits a high density of glucocorticoid receptor expression during early development, exceeding that of the hippocampus (Pavlík and Burešová, 1984). In addition, cerebellar development, including the migration, growth, and maturation of neurons, continues after birth, up to 3 weeks postnatally in rodents and one year postnatally in humans (Ten Donkelaar and Lammens, 2009). Therefore, stressful events occurring during the prenatal period or the first postnatal weeks may adversely affect cerebellar development. Indeed, smaller cerebellar lobe volumes and reduced functional activity of the cerebellar vermis have been identified in children with early deprivation or stress experiences (Bauer et al., 2009).

While classically associated with motor control and coordination, the cerebellum is now increasingly recognized for its involvement in numerous cognitive and emotional functions (Schmahmann, 1998; Adamaszek et al., 2022; Stanley et al., 2023). Accumulating evidence demonstrates its role in language processing, visuospatial function, attention, fear memory, and social behavior through its connections with various brain areas, including the parietal, temporal, and prefrontal cortices, the cingulate gyrus, and the parahippocampal region (Blatt et al., 2013; Badura et al., 2018). Furthermore, many studies highlight the possible implication of the cerebellum in certain psychological disorders, such as autism spectrum disorders, attention deficit hyperactivity disorder, schizophrenia, and anxiety disorders) (Tsai et al., 2012; Yenkoyan et al., 2024). Nevertheless, the effects of early-life stress on the cerebellum have been less explored compared to some other brain areas, including the hippocampus and amygdala.

Here, we examined how MS affects cerebellar circuit maturation and whether these changes are accompanied by alterations in synaptic function and behavior. By integrating molecular, cellular, functional, and behavioral analyses, we identified coordinated changes across multiple levels of cerebellar organization. These findings support the cerebellum as a developmentally sensitive target of early-life adversity and suggest that altered cerebellar maturation may contribute, within broader neural networks, to emotional and social dysfunction.

## 2. METHODS

See Supplemental Methods in Supplement 1 for extended details.

### 2.1 Animals

All experimental procedures were approved by the Italian Ministry of Health (protocol no. 598/2022-PR) and by the Columbia University Institutional Animal Care and Use Committee (AABI2605). Male and female CD1 mice were used for all experiments. Animals were maintained on a 12-hour light/dark cycle with ad libitum access to food and water, at room temperature of 24 °C.

### 2.2 Maternal separation

Pregnant mice were randomly assigned to the control or MS group. Litter sizes were not standardized; all litters were maintained at their natural size. Dams and their litters were housed in home cages containing wood-based bedding and paper-based nesting material. Pups in the MS group were separated from their dams from postnatal day (P)2 to P18 for 4 hours per day (Fig. 1A). During each separation session, pups were kept together as a litter in a separate cage, while dams were housed on another rack. After each session, pups were returned to their home cage and reunited with their dam. Control pups remained undisturbed with their dams until weaning. Following weaning, mice were housed in groups of four with standard environmental enrichment. Mice from each litter were randomly assigned for behavioral, electrophysiological, histological, or molecular analysis at P20, P45, or P60, as indicated in Supplementary Table S1. At P20, no significant difference in body weight was observed between MS and control mice (Supplementary Figure S1).

**Figure 1.**
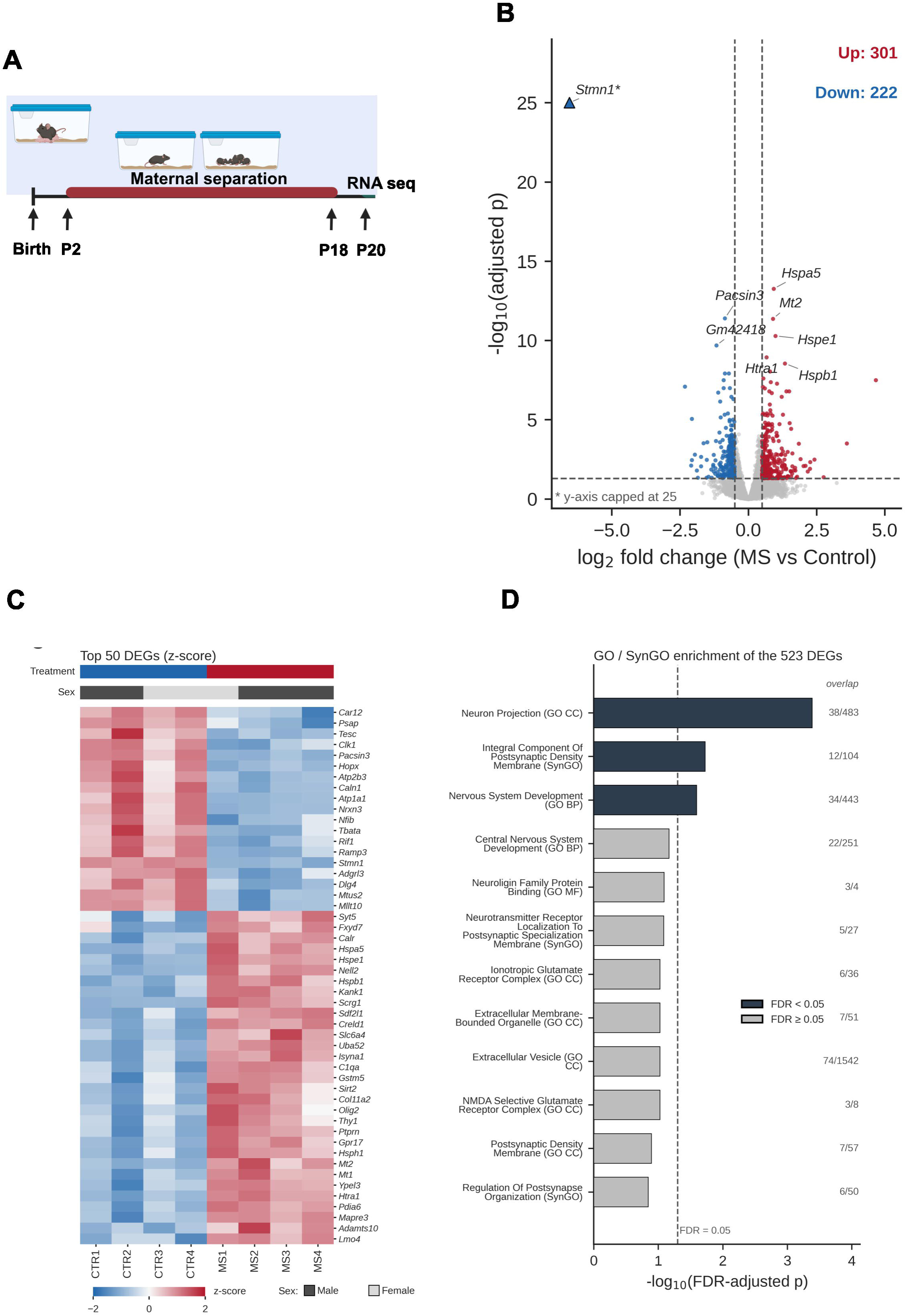
Transcriptional changes in the cerebellum of maternally separated mice. **(A)** Schematic of the maternal separation (MS) protocol (P2–P18, 4 h per day) and of the time point of RNA-seq (P20). **(B)** Volcano plot of 12,289 genes with an adjusted p-value (DESeq2, MS vs Control). Dashed lines: adjusted p = 0.05 and |log2FC| = 0.5. Red: 301 up-regulated DEGs; blue: 222 down-regulated DEGs; gray: not significant. ∗ Stmn1 (adjusted p ≈ 10) is plotted at the cap of the y-axis (25). **(C)** Heatmap of the 50 DEGs with the lowest adjusted p-values (row z-score of VST values); treatment and sex are indicated above the columns. **(D)** GO and SynGO enrichment of the 523 DEGs (Enrichr). Bars show −log10(FDR-adjusted p); dark bars, FDR < 0.05; gray bars, FDR ≥ 0.05; dashed line, FDR = 0.05. Numbers on the right indicate genes overlapping the term/genes in the term. n = 4 animals per group.

### 2.3 RNA sequencing of cerebellar tissues

Total RNA was isolated from the whole cerebellum of P20 male and female mice using the miRNeasy Mini Kit (Qiagen), following the manufacturer’s instructions. RNA quantification was performed using the Qubit 4.0 Fluorometric Assay (Thermo Fisher Scientific). Only the RNA samples that have passed the quality control step with RNA integrity number score of 7.0 or above were used. Libraries were prepared from 125 ng of total RNA using the NEGEDIA Digital mRNA-seq research-grade sequencing service v2.0 (Next Generation Diagnostic srl), which included library preparation, quality assessment, and sequencing on an Illumina NovaSeq 6000 platform using a single-end, 100-cycle strategy. Across the eight samples, the mean sequencing depth was 5.80 million reads per sample, with an average of 78.32% uniquely mapped reads. The raw count data generated in this study are available in Zenodo (Miniaci, 2026).

Differential expression between maternally separated (MS) and control cerebella was tested with DESeq2 (version 1.52.0) on raw counts (MS vs Control; n = 4 per group). P-values were adjusted for multiple testing with the Benjamini–Hochberg procedure (false discovery rate, FDR). Of the 15,246 genes tested, 12,289 had a non-missing adjusted p-value (the remaining genes were excluded by DESeq2 independent filtering or outlier detection). Differentially expressed genes (DEGs) were defined as genes with an adjusted p-value < 0.05 and |log2 fold change| > 0.5, which yielded 523 DEGs (301 up-regulated and 222 down-regulated in MS). Unadjusted p-values are reported for reference only and were not used to define DEGs.

For principal component analysis (PCA), the heatmap and the sex-adjusted model, genes with fewer than 10 total reads across samples were removed (14,208 genes retained) and a blind variance-stabilizing transformation (VST) was applied. PCA was computed on the 500 most variable genes. Sample sex was determined from the expression of Xist and of the Y-linked genes Ddx3y, Uty, Kdm5d and Eif2s3y.

To test whether sex influenced the results, differential expression was repeated with sex as a covariate (design: ∼ sex + condition) using PyDESeq2 0.5.4, applying the same adjusted p-value and fold-change thresholds.

Gene Ontology (Biological Process, Cellular Component, Molecular Function) and SynGO enrichment of the 523 DEGs was performed with Enrichr; terms were considered significant at Benjamini–Hochberg adjusted p< 0.05 (Gene set enrichment analysis (GSEA) was performed with fgsea version on genes ranked by the DESeq2 statistic, using the MSigDB HALLMARK collection (v20255.1Mm), both on protein-coding genes only and on all genes with a non-zero total count; gene sets with adjusted p < 0.05 were considered significant.

The heatmap shows row z-scores of VST values for the 50 DEGs with the lowest adjusted p-values (DEGs absent from the filtered count matrix were skipped). In the volcano plot the y-axis is capped at −log10(adjusted p) = 25.

### 2.4 DNA methylation analysis

Genomic DNA extracted from frozen mouse cerebellum was bisulfite treated and profiled with the Infinium Mouse Methylation BeadChip (Illumina; approximately 285,000 targeted CpGs per sample; Cuomo et al., 2023). IDAT files from six control and six MS mice were processed in RnBeads. Probes overlapping annotated SNPs and probes with unreliable detection measurements (Greedycut detection p-value threshold, 0.05) were removed, followed by exclusion of sex-chromosome probes. No samples were excluded. Methylation data were normalized, leaving 94,006 CpG sites for site-level analysis and principal component analysis. Differential methylation was assessed using limma empirical-Bayes linear models on M-values. P-values were adjusted across the tested sites using the Benjamini-Hochberg false discovery rate method. Effect size was expressed as Δβ = mean βMS − mean βCTRL. Nominal findings were defined by p < 0.05 and FDR-significant findings by adjusted p < 0.05.

### 2.5 Morphology of Purkinje cell dendrites

Purkinje cells were filled with 0.5% biocytin through whole-cell patch-clamp pipette, as previously described (Hoxha et al., 2017), and their dendritic arborization and number of spines were analyzed. To quantify dendritic arbor complexity, we performed Sholl analysis by counting the intersections between dendrites and a series of concentric circles centered on the soma. The initial radius from the center of the soma was 10 μm, and the radial step size was 22 μm, corresponding to sampling radii of 10, 32, 54, 76, 98, and subsequent 22-μm increments. A higher number of intersections across increasing radial distances generally indicates a more extensive and complex dendritic arbor (Sholl, 1953; Bird and Cuntz, 2019). The number of spines/μm of dendritic length is reported as spine density. For more details see supplementary material. PCs were sampled from lobule III and VIII of the cerebellar vermis in both CTRL and MS mice. The same lobules were examined across experimental groups, but sampling was not restricted to a predefined position within the folia. Detailed protocol information is provided in the supplementary materials

### 2.6 Immunostaining

Sagittal cerebellar slices were incubated in blocking buffer (0.5% Triton X-100, 0.2 % bovine serum albumin, 10% goat serum in phosphate-buffered saline) for 4 hours at room temperature. Slices were incubated with primary antibodies: anti-calbindin (mouse; 1:500 dilution), anti-vGLUT1 (rabbit; 1:500 dilution), or anti-vGLUT2 (guinea pig; 1:1000) at 4°C overnight, followed by fluorescent-conjugated secondary antibodies for 2 hours at room temperature. Detailed information on the protocol was provided in the supplementary materials, including the primary and secondary antibodies in Supplementary Table S2. Images were obtained using a Zeiss LSM 900 confocal microscope (Carl Zeiss Microscopy GmbH, Germany).

### 2.7 Electrophysiological recordings

Cerebellar slices were prepared as previously described (Hoxha et al., 2013; Lippiello et al., 2020a, 2020b). Each slice was placed in a recording chamber and perfused with oxygenated artificial cerebrospinal fluid. Whole-cell patch-clamp recordings from PCs were performed by an EPC-8 or EPC-10 patch-clamp amplifier (HEKA Elektronik, Lambrecht/Pfalz, Germany) or MultiClamp 700B amplifier (Scientifica Ltd, UK). Parallel or climbing fibers were stimulated using an electrode placed in the molecular or granular layer, respectively (Binda et al., 2016; Montarolo et al., 2025). Detailed protocol information is provided in the supplementary materials.

### 2.8 RNA extraction, reverse transcription, and qPCR

Total RNA was extracted from the cerebellum of P20 control and MS mice using the Eurogold trifast, and its yield and integrity were assessed spectrophotometrically and by agarose gel electrophoresis. One microgram of RNA was reverse transcribed with LunaScript® RT SuperMix, and qPCR was performed using Luna® Universal qPCR Master Mix on a Bio-Rad CFX96 instrument. Reactions contained cDNA equivalent to 25 ng of RNA and 0.5 μM oligonucleotides. Gene expression was normalized to hydroxymethylbilane synthase (HMBS) and quantified using the 2^-ΔCT^ method. All analyses included at least three biological replicates processed independently. The primers (5’-3’) used for qRT-PCR are listed in Supplementary Table S3.

### 2.9 Western Blot analysis

Cerebellar tissue from P20 control and MS mice was lysed in RIPA buffer (50 mM Tris–HCl pH 8.8, 150 mM NaCl, 0.1% SDS, 0.5% NP-40, 0.5% DOC) with protease and phosphatase inhibitors (ThermoFisher). Lysates were incubated on ice for 30 minutes and clarified by centrifugation at 8,000 × g for 10 min. 20 μg of control and MS lysates were separated by SDS-PAGE and transferred to nitrocellulose or PVDF membranes. Membranes were incubated with the following primary antibodies: BDNF (1:1000, ThermoFisher); CREB (1:1000), p-ERK1/2 (1:1000), ERK1/2 (1:1000), GAPDH (1:1000). After primary antibody incubation, membranes were incubated with HRP-conjugated secondary antibodies (goat anti-rabbit IgG, Amersham, Milan, Italy, 1:10,000) and visualized using chemiluminescence (enhanced chemiluminescence; Euroclone, Milan, Italy). Target protein band intensity was quantified by Quantity One software (Biorad, Milan, Italy) and normalized to GAPDH. Detailed information on the primary and secondary antibodies is provided in supplementary Table S2.

### 2.10 Behavioral Assays

Behavioral assessments at P20, P45, and P60 were performed in independent cohorts, with different animals tested at each age.

#### Beam walk test

The task assessed the ability of mice to walk across an elevated 80 cm-long wood beam (1.5 cm wide). During pre-training, each animal was allowed to traverse the beam twice. Mice were then trained for 3 consecutive days to cross the beam 5 times. The number of footslips, defined as the foot coming off the beam, was measured.

#### Social approach test

Social approach behavior was tested in a three-chambered apparatus, as previously described (Cristiano et al., 2022). The test consisted of 5-min habituation in the empty apparatus, followed by a 10-min test in which one chamber contained a novel sex, strain, and age-matched mouse placed under an enclosed cup, while the other chamber contained an empty cup. Both sides were alternated between the left and right chambers across subjects. The time spent sniffing a novel mouse or a novel object was measured by ANY-maze video-tracking software.

#### Forced swim test

The forced swim test was performed as described previously (Boccella et al., 2021). Mice were placed in a glass beaker filled with 2.5 L of 25°C warm water and recorded for a 6-min period. The amount of time the mouse spent immobile was measured for each trial using a video tracking software (ANY-MAZE, Stoelting Co., Wood Dale, IL, USA).

#### Open field test

Each mouse was placed in a 40 × 40 cm chamber and allowed to explore freely for 30 min (Stanley et al., 2023). The total distance traveled, the number of entries, and time spent in the center region were recorded and analyzed using ANY-maze software.

#### Footprinting

To obtain footprints, the front and back pads were coated in red and blue colors, respectively. The animals were then allowed to walk across a white sheet placed on the floor of a runway (4 cm wide and 40 cm long). The assay was performed in a single trial. The gait parameters were measured for 3 consecutive strides. The walking speed and difference in body weight were balanced to exclude differences, which might affect gait parameters. To characterize the walking pattern of each mouse, we measured the distance between subsequent limb placements (stride length), the overlap between forelimbs and hindlimbs, the distance between the left and right limbs (stance width), and the distance between the right and left hindlimb strides (sway distance).

### 2.11 Drugs

The drugs used for electrophysiology were purchased from HelloBio (Bristol, UK).

### 2.12 Statistics

The Shapiro–Wilk test was first applied to test for a normal distribution of the data. An unpaired Student’s *t*-test or two-way analysis of variance followed by post-hoc multiple comparisons test (GraphPad Prism 9; GraphPad Software) was used for normally distributed data. For data not normally distributed, the Mann-Whitney U test (to compare two groups) and the Kruskal-Wallis test (for multiple-group comparisons) were used. P < 0.05 was considered statistically significant. All data are plotted as mean ± SEM. For all experiments, data from male and female mice were analyzed together. The numbers of males and females in each experimental group are reported in the Supplementary section.

## 3. RESULTS

### 3.1 Transcriptomic analysis of whole cerebellar RNA

To determine how gene expression is affected by MS, bulk RNA-sequencing was performed on whole cerebellar tissue from P20 MS and control mice (Figure 1A). Principal component analysis separated MS from control cerebella along PC1 (62.5% of the variance), whereas PC2 (17.5%) separated the samples by sex, as confirmed by Xist and Y-linked gene expression (Supplementary Figure S2). DESeq2 identified 931 genes with an FDR-adjusted p < 0.05; applying a fold-change threshold (|log2FC| > 0.5), we defined 523 DEGs (301 up-and 222 down-regulated in MS (Figure 1B-C; Supplementary Table S4). Among the most significant were the down-regulated Stmn1, Pacsin3 and Atp2b3, and the up-regulated Hspa5, Mt1/Mt2, Hspe1 and Hspb1.

Enrichment analysis of the 523 DEGs showed that, after FDR correction, three terms were significantly enriched: Neuron Projection (GO Cellular Component; 38/483 genes, adjusted p = 4.1 × 10), Nervous System Development (GO Biological Process; 34/443, adjusted p = 0.025) and Integral Component of Postsynaptic Density Membrane (SynGO; 12/104, adjusted p = 0.019) (Figure 1D; Supplementary Table S6). Other synaptic terms (for example, the NMDA and ionotropic glutamate receptor complexes and neuroligin binding) showed only nominal enrichment that did not survive FDR correction and should be regarded as suggestive.

GSEA of the HALLMARK collection showed a consistent signal for four gene sets, significant both in protein-coding genes and in all genes: positive enrichment of mTORC1 signaling and negative enrichment of the G2M checkpoint. Five further sets (positive: Notch signaling, PI3K/AKT/mTOR signaling, Fatty Acid Metabolism, Complement; negative: E2F targets) reached significance only when the analysis was restricted to protein-coding genes (adjusted p 0.033– 0.048; 0.058–0.063 in all genes) and are therefore considered suggestive (Supplementary Figure S3; Supplementary Table S7).

Because sex explained a substantial fraction of variance, we repeated the analysis with sex as a covariate. Log2 fold changes were highly correlated with those of the main analysis (r = 0.97), and 432 of the 523 DEGs were also detected (561 DEGs in total; Supplementary Figure S4A-B; Supplementary Table S5). Among the genes with a significant sex effect, four of five were Xist or Y-linked genes (Supplementary Figure S4C).

Overall, these data indicate that early-life MS alters cerebellar transcriptional programs, including genes involved in neuronal development, neuronal projections, and postsynaptic organization.

### 3.2 Ultrastructural modifications of Purkinje cell dendrites in maternally separated mice

Given the enrichment of transcripts related to neuronal architecture and synaptic remodeling, we next examined whether MS induces structural alterations in cerebellar neurons (Figure 2A). We, therefore, examined the dendritic arborization of Purkinje cells (PCs) in MS mice using Sholl analysis. At P20, PCs from MS mice displayed an increased number of intersections in the proximal region of the dendritic arbor compared to controls, especially at 54 and 76 μm from the soma (Figure 2B). However, the number of dendritic spines was comparable between MS and control PCs (Supplementary Figure S5).

**Figure 2.**
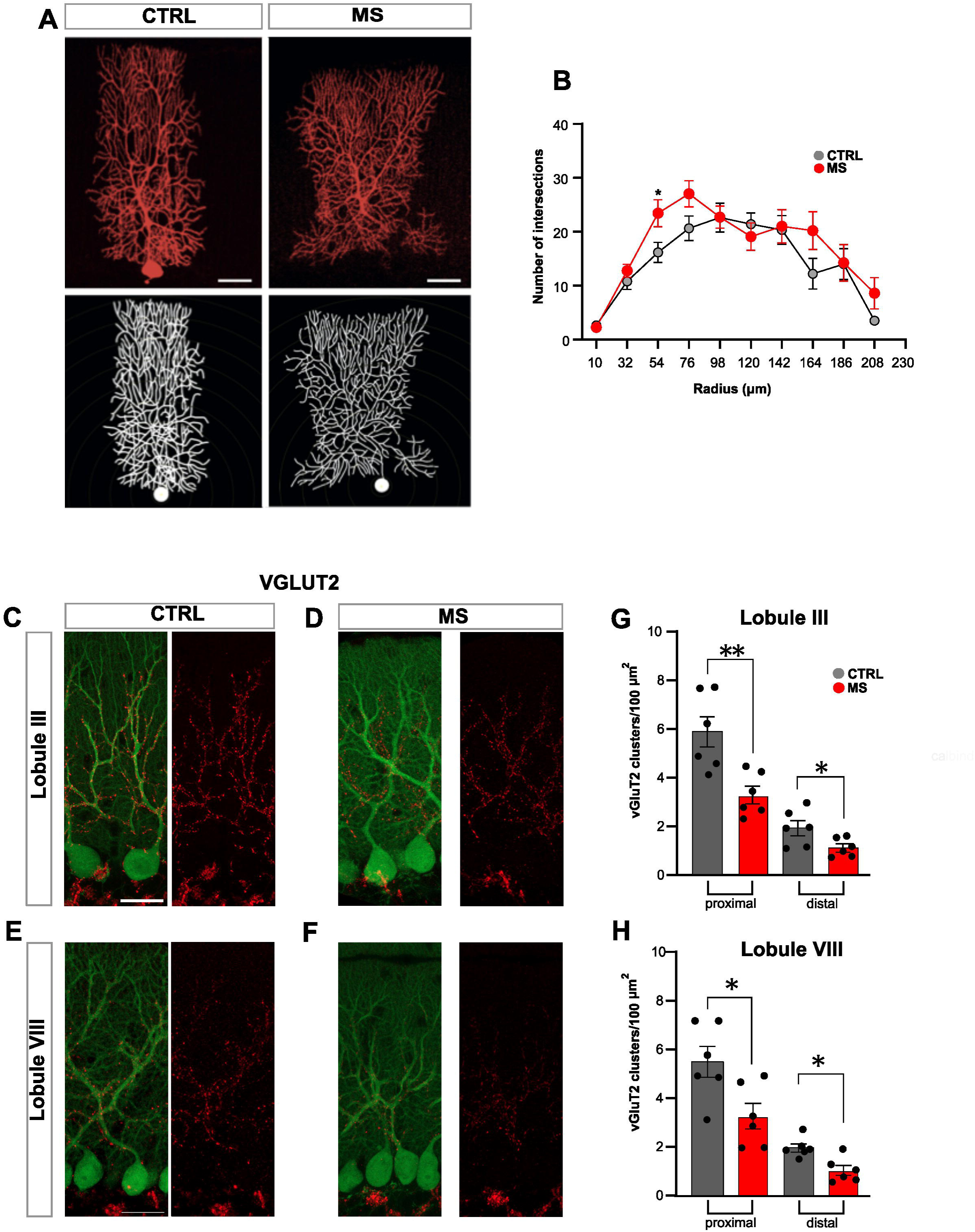
Purkinje cell dendritic tree extension in maternal separated mice. **(A)** Biocytin-filled Purkinje cells (PCs) and reconstructed structure for control (CTRL) and maternal-separated (MS) mice at P20. Scale bar 30 μm. **(B)** Sholl analysis showing the number of dendritic intersections as a function of distance from the soma (10–208 µm) for MS PCs (n = 11 cells) compared to their control littermates (n= 8 cells; n = 4-5 mice per group). Data were analyzed using a two-way mixed-effects model, accounting for repeated measurements across radial distances and matching by litter with treatment as the fixed effect and litter as the random effect. For the full radial range: Treatment effect (F_(1,4)_ = 0.91, p =.40); radius effect (F_(9,33)_ = 28.18, p = 0.0001); radius by treatment interaction (F_(9,33)_ = 1.08, p = 0.40); for CTRL (n = 11 cells/4 mice), for MS (n = 18 cells/5 mice). For the proximal region (distances 10-98 µm from the soma): treatment effect (F_(1,4)_ = 2.24, p = 0.21); radius effect (F_(4,16)_ = 111.6, p = 0.0001); radius by treatment interaction (F_(4,16)_ = 4.29, p = 0.02); Bonferroni-adjusted post hoc comparison at 54 µm: t = 5.12, df = 3.52, adjusted p = 0.040. **(C, D, E, F)** Representative confocal images of CTRL and MS mice at P20 of cerebellar lobule III and VIII sections, double-labeled for calbindin (green, Purkinje cells) and vGluT2 (red, climbing-fiber synapses). Scale bar 25 µm. (**G, H**) Quantitative analysis revealed a lower density of VGLUT-2 puncta in the proximal and distal dendritic compartments of PCs of MS mice than controls in lobule III (proximal t_10_=3.605, p= 0.004; distal t_10_=2.255, p= 0.047) and VIII (proximal t_10_=2.706, p= 0.022; distal t_10_=2.252, p= 0.048). Each dot represents data from an individual mouse; for each mouse, three sections were analyzed, with four image fields acquired per section (n = 6 mice per group). Unpaired t-test analysis. ∗p <.05, ∗∗p <.01.

Since the proximal dendritic compartment of Purkinje cells is the main territory of climbing-fiber innervation, we next analyzed the expression of the vesicular glutamate transporter VGLUT2, a marker of climbing-fiber terminals (Fremeau et al., 2001). Lobules III and VIII were selected as representative regions of the anterior and posterior cerebellum (Ozol et al., 1999) to assess regional differences along the rostrocaudal axis.

Quantification in cerebellar lobules III and VIII revealed a significant reduction in the density of VGLUT2-positive puncta in MS mice compared with controls (Figure 2C–H; Supplementary Figure S6). By contrast, VGLUT1 expression, which mainly labels parallel-fiber terminals in the molecular layer, was not significantly altered in MS mice (Supplementary Figure S7).

Together, these findings indicate that MS alters the structural organization of PCs, characterized by increased proximal dendritic complexity but reduced density of VGLUT2-positive puncta. This suggests an abnormal remodeling of excitatory cerebellar connectivity following early-life stress.

### 3.3 Cerebellar synaptic plasticity is impaired in mice experiencing maternal separation

We next examined whether these structural alterations were accompanied by changes in cerebellar synaptic function. Patch-clamp recordings were performed in acute cerebellar slices to assess synaptic transmission and plasticity at the parallel fiber (PF)-PC synapse, which is considered the main plastic site of the cerebellar cortex. Analysis of spontaneous excitatory postsynaptic currents (Supplementary Figure S8) and evoked EPSC amplitudes in response to varying PF stimulation intensities showed no significant differences between MS and control mice, suggesting that the basal PF-PC synaptic transmission was not affected by early-life stress (Supplementary Figure S9A). Moreover, paired-pulse facilitation at the PF-PC synapse was comparable between groups across the tested interstimulus intervals, suggesting that short-term plasticity at this synapse was also not affected by MS (Supplementary Figure S9B). In contrast, paired-pulse stimulation at the climbing fiber (CF)-PC synapse revealed a significantly reduced short-term depression in MS mice at the interstimulus interval of 50 ms (Supplementary Figure S10), indicating a selective alteration of CF–PC synaptic function.

We next examined the two main forms of cerebellar synaptic plasticity, long-term potentiation (LTP) and long-term depression (LTD) at PF-PC synapses. As shown in Figure 3A-B, tetanic stimulation of PFs elicited LTP in PCs of control mice, while no potentiation was observed in MS slices taken at different ages, from P20 to P60. LTD was successfully induced in control slices following PF burst stimulation in conjunction with PC depolarization (Figure 3C-D). In contrast, a significant deficit in LTD was observed in MS mice at P20, whereas at P45 and P60, LTD levels did not differ from those of controls. These findings indicate that MS produces persistent impairment of PF-PC LTP, together with an early deficit in LTD.

**Figure 3.**
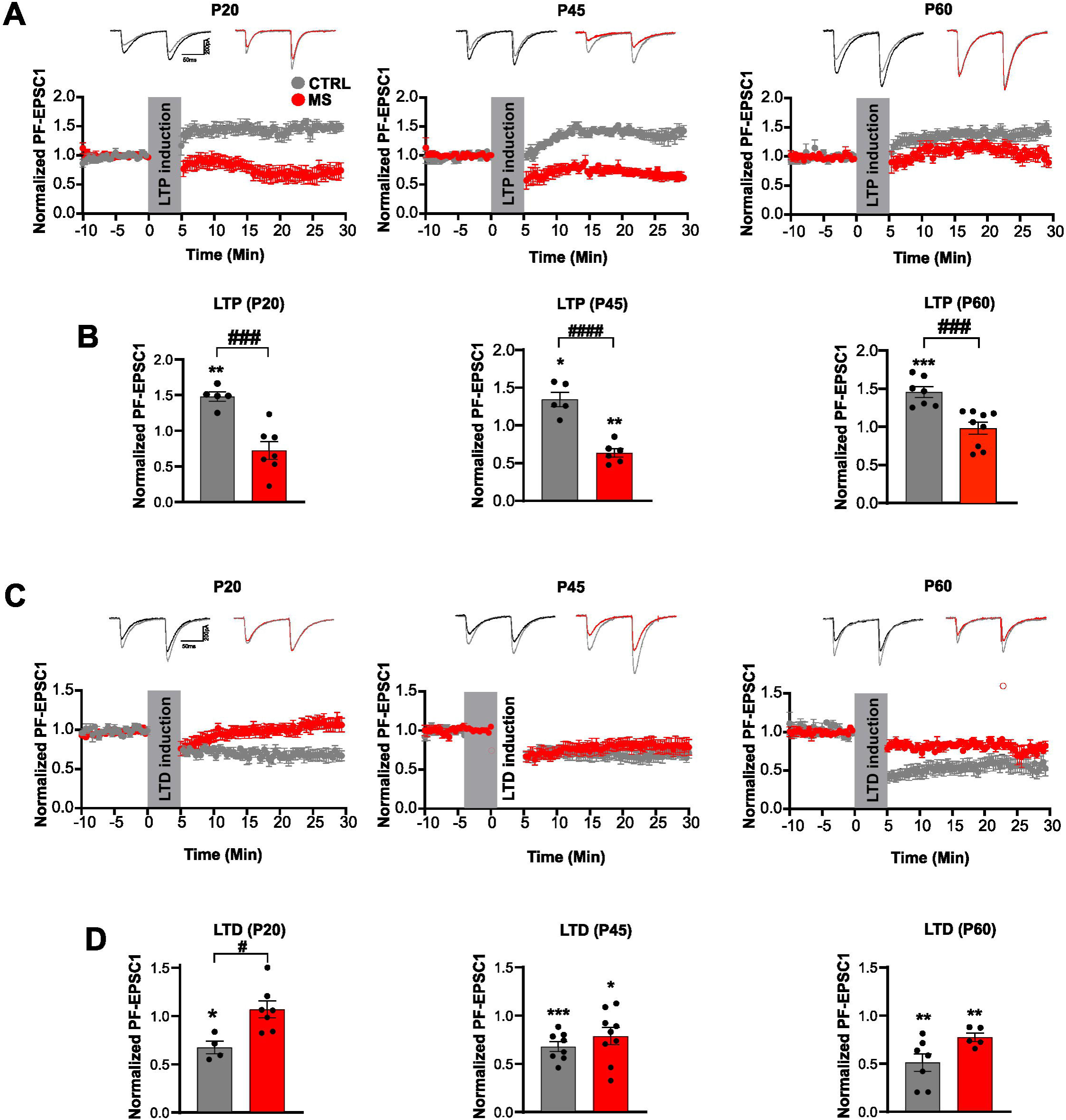
Cerebellar synaptic plasticity is impaired in maternal-separated mice. **(A)** Time course of normalized parallel fiber (PF)-EPSC1 amplitude before and after long-term potentiation (LTP) induction in cerebellar slices of MS and control mice (n =5-9 cells per group; n= 5-6 mice per group). The top inserts show superimposed representative recordings of PF-EPSCs evoked by paired-pulse stimulation before and 30 min after tetanic stimulation. **(B)** Normalized PF-EPSC1 recorded 30 min after LTP induction in control (vs baseline at P20 t_4_=6.673, p= 0.002; at P45 t_4_=3.676, p= 0.021; at P60 t_6_=6.992, p= 0.0004) and MS mice (vs baseline at P20 t_6_=2.315, p= 0.059; at P45 t_5_=6.587, p= 0.001; at P60 t_8_= 0.857, p= 0.85). LTP deficits were observed in MS mice at each age compared to control mice (at P20 t_10_=4.798, p= 0.0007; at P45 t_9_=6.699, p<.0001; at P60 t_14_=4.623, p= 0.0004). **(C)** Time course of normalized PF-EPSC1 amplitude before and after long-term depression (LTD) induction in cerebellar slices of MS and control mice (n = 5-9 cells per group; n= 5-7 mice per group). The inserts on the top show superimposed traces evoked before and 30 min after LTD induction. **(D)** Normalized PF-EPSC1 at 30 min after LTD induction in control (vs baseline at P20 t_3_=5.167, p= 0.014; at P45 t_7_=6.239, p= 0.0004; at P60 t_6_=5.382, p= 0.0017) and MS mice (vs baseline at P20 t_6_=0.925, p= 0.39; at P45 t_8_=2.426, p= 0.041; at P60 t_4_=5.153, p= 0.0067). LTD impairment was observed in MS mice at P20 vs control (at P20 t_9_=3.076, p= 0.013; at P45 t_15_=1.025, p= 0.32; at P60 t_10_=2.274, p= 0.05). Each dot represents data from an individual Purkinje cell. Paired and unpaired t-test analysis. ∗*p <*.05, ∗∗*p <*.01, ∗∗∗*p <*.001 vs baseline; #*p <*.05, ##*p <*.01, ###p <.001; #### *p <*.0001 MS vs control mice.

### 3.4 Effect of maternal separation on the cerebellar BDNF signaling pathway

During the critical period of brain development, the BDNF signaling pathway plays a crucial role in forming and refining neuronal circuits by promoting synapse formation and plasticity (Colucci-D’Amato et al., 2020). Since BDNF expression is dysregulated by stress and is implicated in the pathophysiology of mood disorders (Castrén and Monteggia, 2021; Sun et al., 2023), we examined the effect of MS on cerebellar BDNF expression at the transcriptional and translational levels via RT-qPCR and Western blotting, respectively. We found that both the BDNF transcript and protein levels were significantly reduced in MS mice at P20 compared to controls (Figure 4A-C).

**Figure 4.**
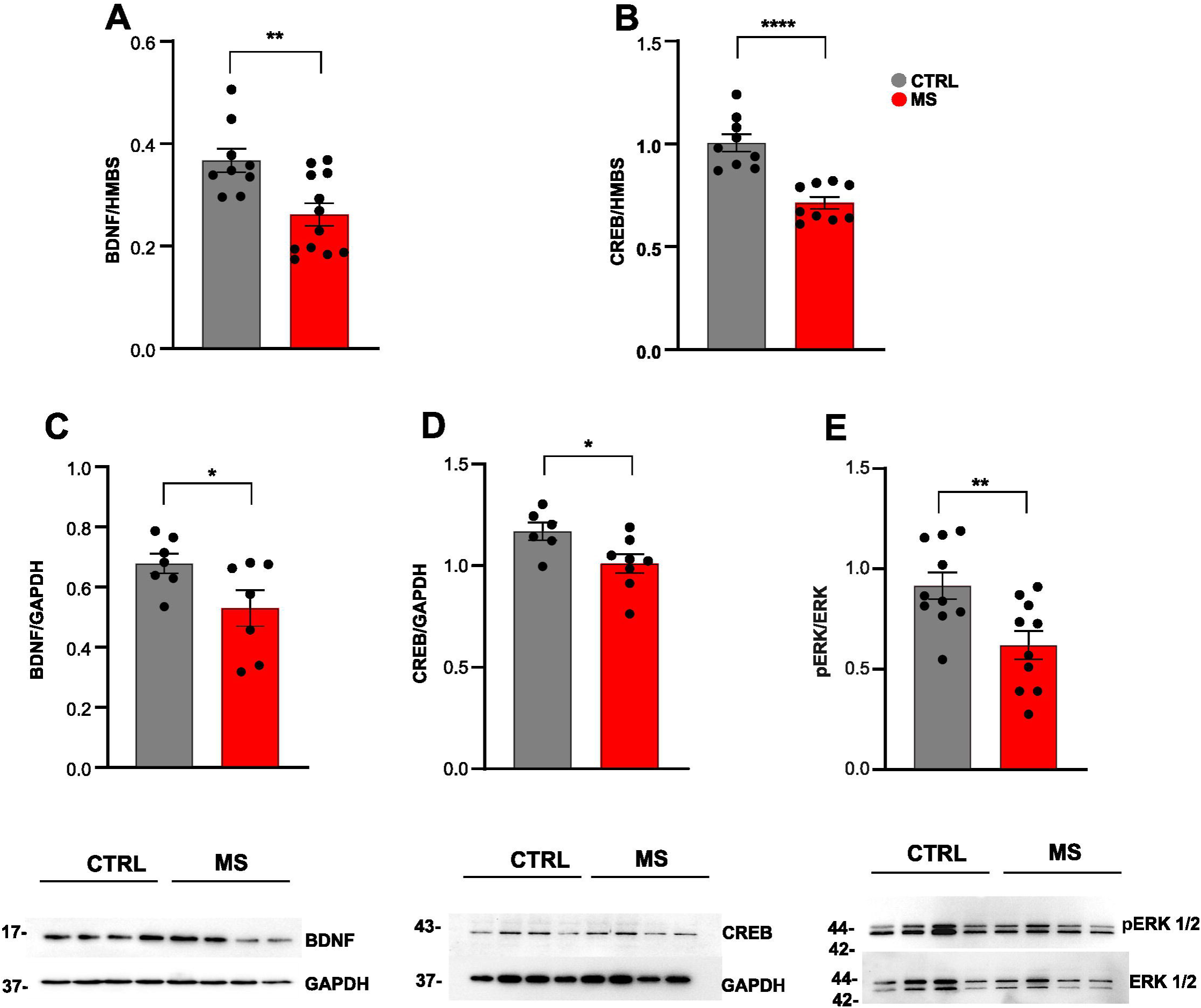
Maternal separation alters BDNF and CREB signaling pathways in the offspring mouse cerebellum. (A, **B)** BDNF and CREB mRNA expression levels in duplicate in maternal separated (MS) and control (CTRL) mice at P20 (for BDNF, t_19_ = 3.261, df_19_, *p*=0.041; for CREB, t_16_ = 5.691, df_16_, *p<*.0001). Gene expression levels were normalized to hydroxymethylbilane synthase (HMBS). **(C, D)** Western blot analysis revealed a significant reduction of BDNF, CREB, and pERK in MS compared to control mice (for BDNF t_12_ =2.182, p= 0.049; for CREB t_12_ =2.417, p= 0.032; for pERK t_18_ =3.042, p= 0.007). Protein levels of BDNF and CREB in control and maternal mice were normalized to GAPDH. The level of ERK phosphorylation was normalized to unphosphorylated ERK (ERK1/2). The insets at the bottom display representative western blots probed with antibodies against BDNF, CREB, GAPDH, p-ERK, and ERK. The molecular weight of each protein is indicated on the left in kDa. Each dot represents data from an individual mouse (n= 6-10 mice per group). Unpaired t-test analysis. ∗*p <*.05, ∗∗*p <*.01, ∗∗∗*p <*.001.

Since BDNF acts through the CREB and ERK pathways to regulate synaptic plasticity (Lado et al., 2025), we also assessed the expression levels of these two key proteins. Compared to the control group, P20 MS mice showed a marked reduction in CREB mRNA and protein expression (Figure 4B-D), along with a significant decrease in ERK phosphorylation levels (Figure 4E).

These findings indicate that MS is associated with reduced BDNF-related signaling in the cerebellum.

### 3.5 Behavioral effects of maternal separation in the offspring

To determine whether MS was associated with behavioral alterations in the offspring, we performed a battery of tests assessing motor, social, and stress-coping phenotypes. In the beam walking test, MS mice showed an increased number of footslips, indicating impaired balance and motor coordination (Figure 5A). However, footprinting analysis (Supplementary Figure S11) and open-field analysis (Figure 5B) did not reveal abnormalities in gait pattern or spontaneous locomotor behavior, suggesting that the impaired beam-walking performance was not associated with a generalized motor impairment.

**Figure 5.**
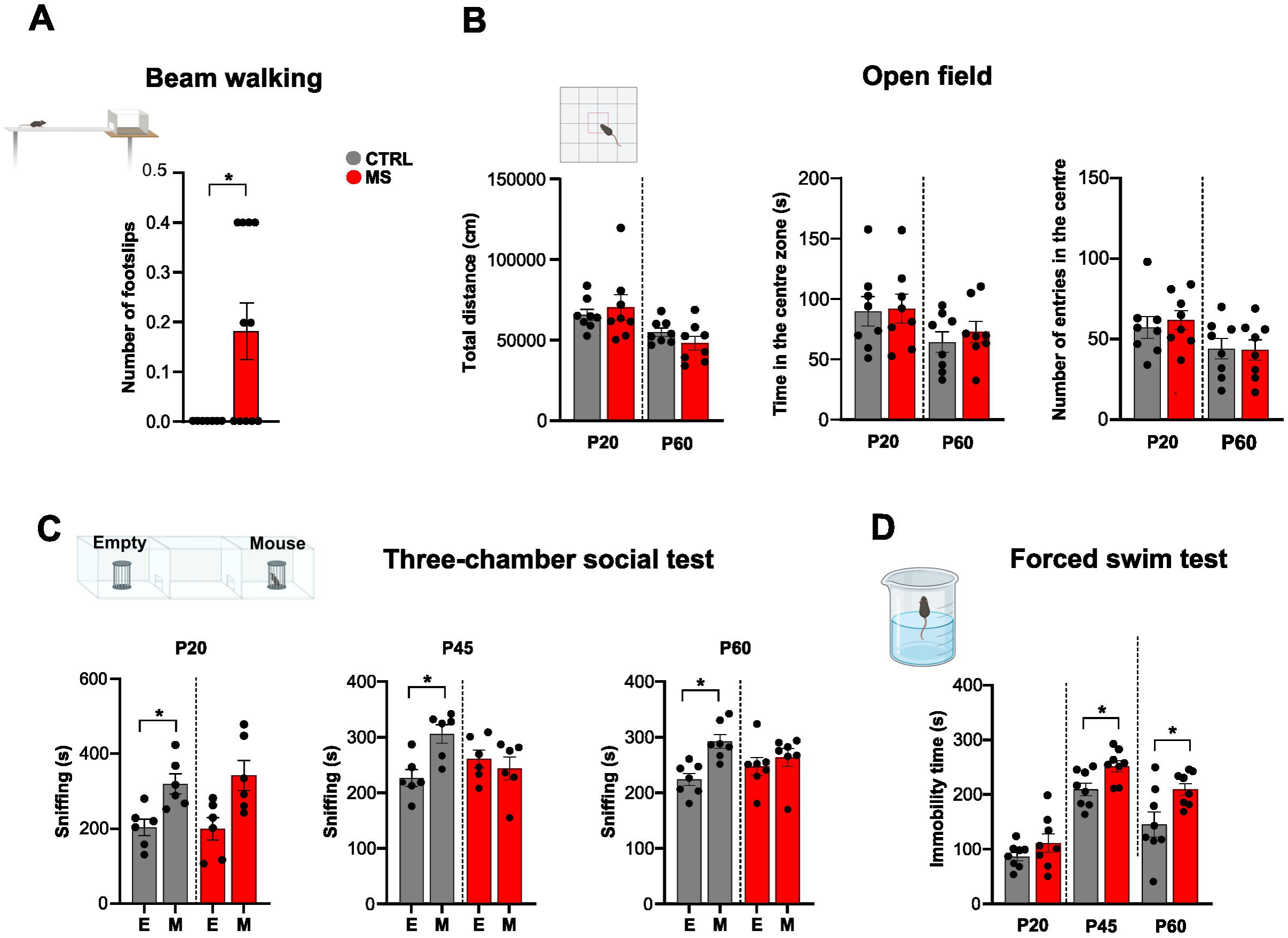
Effect of maternal separation on offspring behavior. **(A)** P60 MS mice exhibited a higher number of footslips than controls when crossing the 1.5-cm-wide beam (t_16_=2.520, p= 0.022). **(B)** The open field analysis revealed no difference between MS and control mice regarding total distance traveled (at P20 t_14_=0.494, p= 0.628; at P60 t_14_= 1.386, p= 0.187), time (at P20 t_14_=0.131, p= 0.896; at P60 t_14_=0.686, p= 0.503), and number of entries in the center (at P20 t_14_=0.503, p= 0.622; at P60 t_14_=0.84, p= 0.934). **(C)** In the three-chamber test, MS mice did not show any preference for either the empty (E) or the mouse (M) chamber at all ages tested (at P20 t_5_=2.053, p= 0.095; at P45 t_5_=0.491, p= 0.643; at P60 t_6_=0.922, p= 0.391), while the control mice spent more time with the other mouse (at P20 t_5_=2.638, p= 0.046; at P45 t_5_=2.663, p= 0.044; at P60 t_6_=3.106, p= 0.021). **(D)** In the forced swim test, maternal separated (MS) mice (red bars) displayed more immobility time at P45 (t_14_=2.722, p= 0.165) and P60 (t_14_=2.537, p= 0.023) compared to control (CTRL) mice (grey bars). Each dot represents data from an individual mouse (n= 6-11 mice per group). Paired or unpaired t-test analysis. ∗*p <*.05, ∗∗*p <*.01, ∗∗∗∗*p <*.0001.

The motor phenotype was accompanied by altered social behavior in the three-chamber test: whereas control mice consistently preferred the novel mouse over the empty cup, MS mice showed no clear social preference across the ages tested, suggesting reduced sociability (Figure 5C). MS mice also displayed increased immobility in the forced swim test at P45 and P60, indicating the emergence of altered stress-coping behavior during adolescence and adulthood (Figure 5D). Overall, these findings indicate that MS produces a selective behavioral phenotype, characterized by impaired motor coordination together with concurrent alterations in social approach and stress-coping behavior.

### 3.6 DNA methylation and expression of glucocorticoid receptors

Given the central role of glucocorticoid receptor signaling in stress responses and developmental plasticity, we next examined whether MS affects glucocorticoid receptor methylation and expression in the cerebellum. Indeed, several studies indicate that early-life stress induces epigenetic modifications at the glucocorticoid receptor promoter, leading to decreased receptor expression and, consequently, heightened HPA axis responsiveness to stress (Ladd et al., 2004; Navailles et al., 2010). We, therefore, performed DNA methylation profiling combined with analysis of glucocorticoid receptor mRNA expression in MS mice and controls. The DNA methylation array covered 194,006 CpG sites, of which 20,503 were differentially methylated between MS and control mice at P20 (Figure 6A). Interestingly, a CpG site in the glucocorticoid receptor (GR) promoter was hypermethylated in cerebellar tissue from MS mice compared with controls (Figure 6B). RT-qPCR (Figure 6C) and Western blot analyses (Figure 6D) further showed reduced GR mRNA and protein levels at P20 (Figure 6C–D), alongside increased promoter methylation. Together, these findings indicate that maternal separation is associated with GR promoter hypermethylation and reduced GR expression in the developing cerebellum.

**Figure 6.**
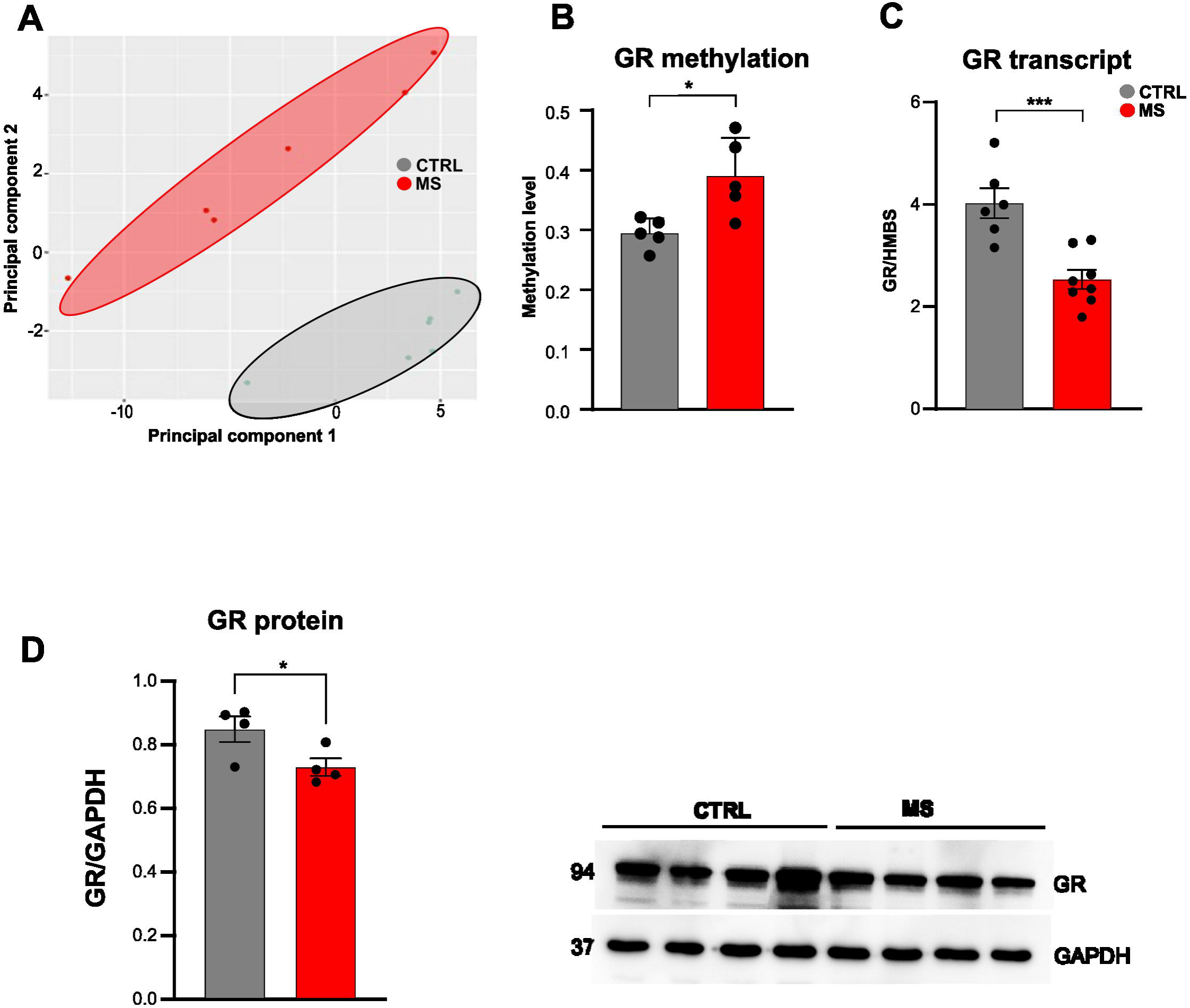
Maternal separation affects the methylation of the glucocorticoid receptor promoter. **(A)** Principal component analysis was performed on the methylation state of 285.000 CpG sites for each mouse at P20. PC1 and PC2 explain 55.2% and 13.3% of the total variance, respectively (68.5% cumulative variance). The two clusters indicate that maternal-separated (MS) and control (CTRL) mice have distinct methylation profiles. (**B)**. Methylation levels of the glucocorticoid receptor (GR) promoter (cg35634261_BC21) in MS and control mice (t_8_=3.104, p= 0.014). **(C)** GR mRNA expression in MS and control mice normalized to hydroxymethylbilane synthase (HMBS) (t_12_=4.520, p= 0.0007). (D) Western blot analysis revealed a significant reduction of GR in MS compared to control mice (for GR t_6_ =2.456, p= 0.049). Protein levels of GR in control and maternal mice are normalized to GAPDH. The insets on the bottom display representative western blots probed with antibodies against GR and GAPDH. The molecular weight of each protein is indicated on the left in kDa. Each dot represents data from an individual mouse (n = 4-8 mice per group). Unpaired t-test analysis. ∗*p <*.05, ∗∗∗*p <*.001

## 4. DISCUSSION

This study demonstrates that MS induces long-lasting alterations in cerebellar maturation, identifying a convergent and pathway-selective cerebellar phenotype comprising Purkinje-cell dendritic remodeling, persistent impairment of synaptic plasticity, and reduced neurotrophic signaling.

At the molecular level, RNA-sequencing of whole cerebellar tissue revealed early changes in gene-expression programs related to synaptic transmission, cytoskeletal organization, neuronal architecture, and plasticity. These processes are particularly relevant during early postnatal cerebellar development, when PC dendrites expand, and both climbing-and parallel-fiber synaptic inputs are progressively refined (Hashimoto et al., 2009; Nitta et al., 2025). The concomitant dysregulation of genes involved in transcriptional and epigenetic regulation further suggests that MS affects not only synaptic components but also the regulatory pathways that shape how cerebellar cells respond to stress-related signals.

Consistent with the transcriptional profile, MS induced structural remodeling of PCs, characterized by increased dendritic complexity particularly in the proximal region of the dendritic tree. This structural modification was accompanied by reduced density of presynaptic glutamate transporter VGLUT2, whereas VGLUT1 expression in the molecular layer remained unchanged. Increased dendritic complexity and reduced puncta density describe distinct aspects of remodeling and are not necessarily contradictory. Reduced puncta density may reflect changes in VGLUT2 abundance or the distribution of labeled terminals relative to the dendritic arbor and does not, by itself, establish a loss of climbing-fiber innervation. Our electrophysiological findings provide complementary evidence of altered short-term dynamics at CF–PC synapses but do not demonstrate a reduction in the number of functional climbing-fiber inputs.

The interpretation of VGLUT2 labeling also requires consideration of developmental stage. In the mature cerebellar molecular layer, VGLUT1 and VGLUT2 are commonly used as markers of PFs and CFs, respectively (Hioki et al., 2003; Zander et al., 2010). However, the developmental switch from VGLUT2 to VGLUT1 in PF terminals is not completed until P30. VGLUT2-positive puncta at P20 may therefore include immature PF terminals and cannot therefore be attributed exclusively to CFs (Miyazaki et al., 2003).

The dendritic remodeling observed in our study should also be considered in the context of the region-specific effects of early-life stress on neuronal architecture. For example, MS has been shown to lead to dendritic shrinkage and reduced spine density in the hippocampus and medial prefrontal cortex, while promoting dendritic growth in the basolateral amygdala (Monroy et al., 2010; McEwen et al., 2016). In the cerebellum, Oliveira et al. reported increased Purkinje-cell layer height and cytoplasmic area in maternally separated UChA rats, together with reduced Purkinje-cell perimeter in UChB rats, whereas the molecular and granular layers and white matter were unaffected (Oliveira et al., 2014). Our results show that MS also alters PC dendritic architecture, with increased complexity in the proximal region of the dendritic tree. Whether these dendritic changes reflect compensation for altered synaptic signaling or disrupted developmental refinement remains to be established.

Electrophysiological recordings further support the idea of pathway-specific rather than global synaptic dysfunction. Basal PF–PC transmission was preserved in MS mice, as indicated by unchanged spontaneous EPSCs, input-output responses, and paired-pulse facilitation. This functional preservation is consistent with the unchanged VGLUT1 expression and suggests that PF synaptic transmission remains largely intact under basal conditions. In contrast, paired-pulse depression at CF–PC synapses were reduced at short interstimulus intervals, indicating altered climbing-fiber synaptic dynamics. Because paired-pulse depression at CF–PC synapses depends largely on presynaptic mechanisms (Hashimoto and Kano, 1998), this finding suggests that MS modifies the functional properties of the climbing-fiber input, in agreement with the reduction in VGLUT2 expression and the structural changes observed in the proximal PC compartment.

The most persistent functional alteration induced by MS was the impairment of PF–PC long-term potentiation. Indeed, PF-PC LTP was disrupted from the juvenile to the adult stages, whereas LTD was mainly affected at P20 and recovered later in life. This persistent LTP deficit is particularly relevant because PF–PC synapses represent a central site through which PCs integrate cerebellar inputs and regulate cortical output. PFs convey mossy fiber-derived sensory, motor, and associative information to PCs, while climbing fibers from the inferior olive provide instructive signals that shape the direction of plasticity (Strick et al., 2009; Palesi et al., 2017). Within this circuit, coincident PF and climbing fiber activity favors LTD, whereas repeated PF activation in the absence of climbing fiber input promotes LTP (Coesmans et al., 2004; Lippiello et al., 2020b). Therefore, a sustained impairment of PF–PC LTP may reduce Purkinje cells’ ability to adjust their responsiveness during development and in response to experience. Because PCs provide the sole output of the cerebellar cortex, altered PF–PC plasticity is likely to affect the communication of cerebellar circuits with motor, cognitive, social, and emotional networks.

The mechanisms underlying the impairment of functional synaptic plasticity following MS likely involve interactions among neurotrophic signaling, stress-response pathways, and epigenetic regulation. One such mechanism may be the downregulation of BDNF and CREB that we observed in MS cerebellar tissue, both of which are critical for synaptic maintenance and plasticity. The elevated glucocorticoid levels commonly associated with ELS may further contribute to these impairments. Indeed, stress exposure has been shown to impair hippocampal LTP through glucocorticoid receptor activation and to reduce BDNF expression in the dentate gyrus, effects that can be mimicked by corticosterone administration (Smith et al., 1995). A growing body of evidence points to multifaceted crosstalk between glucocorticoids and BDNF (Tsimpolis et al., 2024): the two systems share intracellular signaling pathways and reciprocally influence each other’s expression and function. Both in vitro and in vivo studies have shown that glucocorticoids suppress the expression of BDNF mRNA and its receptor, TrkB, in hippocampal neurons (Schaaf et al., 1998; Lee et al., 2020). Reduced BDNF protein levels have been reported in the rat hippocampus following chronic corticosterone exposure in parallel with the emergence of depressive-like behavior (Jacobsen and Mørk, 2006; Huang et al., 2011).

Our epigenetic findings further support the involvement of stress-response pathways in cerebellar dysfunction. We found increased methylation of the glucocorticoid receptor promoter together with reduced receptor transcript levels in the cerebellum. However, further studies are needed to establish a direct causal link between promoter methylation and GR expression. Previous studies show that early-life stress can induce epigenetic modifications of the glucocorticoid receptor gene in the hippocampus and cortex, reducing receptor expression and altering HPA-axis feedback (Ladd et al., 2004; Navailles et al., 2010). In the cerebellum, Wilber and collaborators reported a different pattern: neonatal MS increased glucocorticoid receptor immunoreactivity selectively in the posterior interpositus nucleus of adult male rats, correlating with poorer eyeblink conditioning (Wilber et al., 2007). The different direction of the glucocorticoid receptor changes may reflect differences in separation protocol, developmental stage, cerebellar compartment, and molecular readout. In particular, they assessed glucocorticoid receptor protein immunoreactivity in a restricted region of the adult cerebellar nuclei, whereas our results add an epigenetic dimension by showing that glucocorticoid receptor promoter hypermethylation accompanies reduced cerebellar receptor transcription during a critical period of circuit maturation. Region-and cell-type-specific analyses across development will be required to determine how these adaptations evolve and interact with altered neurotrophic signaling, synaptic refinement, and plasticity.

The behavioral findings provide the clearest functional context for the cerebellar phenotype. MS mice made more footslips during beam walking, whereas gait parameters and spontaneous locomotion remained intact. This pattern argues against generalized motor weakness, locomotor suppression, or a basic gait abnormality and instead supports a selective deficit in demanding balance and coordination. Impaired cerebellar-dependent eyeblink conditioning has been demonstrated after neonatal maternal separation, providing independent evidence that early-life stress can have long-lasting consequences for behaviors that depend strongly on cerebellar circuitry (Wilber *et al*, 2007).

MS was also accompanied by altered social investigation and increased forced-swim immobility. These findings are consistent with the broader behavioral consequences of early-life adversity (Waters and Gould, 2022; Takabayashi et al., 2025) but cannot be assigned specifically to cerebellar dysfunction from the present experiments. Social behavior and stress-coping responses engage distributed networks involving prefrontal, limbic, striatal, hypothalamic, and cerebellar regions. The cerebellum may contribute to these phenotypes through its extensive connections with nonmotor circuits, but MS also affects multiple extracerebellar regions that likely contribute to the overall phenotype.

## 5. LIMITATIONS OF THE STUDY

The molecular analyses were performed on whole cerebellar tissue and therefore do not establish whether the observed alterations occur within Purkinje cells or other cerebellar cell populations. Their relationship to Purkinje-cell physiology should consequently be interpreted at the circuit and tissue levels rather than as evidence of a Purkinje cell-specific molecular mechanism. In addition, the present data do not establish whether the molecular changes observed at P20 persist into adulthood or account for the sustained impairment of synaptic plasticity. Both male and female mice were included, but the small number of animals of each sex limits our ability to determine whether maternal separation affects males and females differently. Finally, while control pups remained undisturbed, MS pups experienced both maternal absence and handling. This design does not allow us to distinguish the effects of maternal absence from those of handling or possible changes in maternal care after reunion.

## 6. CONCLUSIONS

Overall, the present study shows that maternal separation selectively remodels Purkinje-cell circuitry, alters climbing-fiber-associated organization, and produces a persistent deficit in long-term synaptic plasticity. These circuit changes emerge together with reduced BDNF-related signaling and altered glucocorticoid receptor regulation and are accompanied by impaired performance on a demanding motor-coordination task. By connecting early molecular and epigenetic alterations with pathway-specific structural and physiological changes across development, this study advances previous observations of cerebellar vulnerability and identifies Purkinje-cell circuit plasticity as a persistent target of early-life adversity.

## DATA AVAILABILITY

The raw data described in this article will be made available by the authors without undue reservation.

## AUTHOR CONTRIBUTIONS

MCM, LC, and DS conceived and designed the experiments. CC, AS, FML, and PL performed behavioral experiments and analysis. PL and SP performed electrophysiological experiments. EH, LS, and AMR performed the structural and immunofluorescence analyses. MGF, SP, and KDF performed molecular biology assays under the supervision of FV. SDA and DT performed the analysis of RNA-sequencing data. MC performed the methylation analysis under the supervision of LC. MCM wrote the paper in consultation with all other authors.

## ACKNOWLEDGEMENTS

We thank Antonio Baiano, Angelo Russo, and Giovanni Esposito for their invaluable contributions in managing the animal facility, which made this study possible.

This work was supported by the Ministry of University and Research (MUR), the National Recovery and Resilience Plan (NRRP), project MNESYS (PE0000006) – A Multiscale integrated approach to the study of the nervous system in health and disease (DN. 1553 11.10.2022), and the Internationalization grants from the University of Naples Federico II to MCM. AS and DS were supported by the Freedom Together Foundation and NIH NIDA R01DA07418. DT was supported in part by a research grant from Research Ireland under Grant Number 21/RC/10294_P2 and co-funded under the European Regional Development Fund and by FutureNeuro industry partners.

## Generative AI statement

The author(s) declared that generative AI was not used in the creation of this manuscript.

