## Supplement 1 for "Maternal separation remodels Purkinje cell circuits and impairs cerebellar synaptic plasticity"

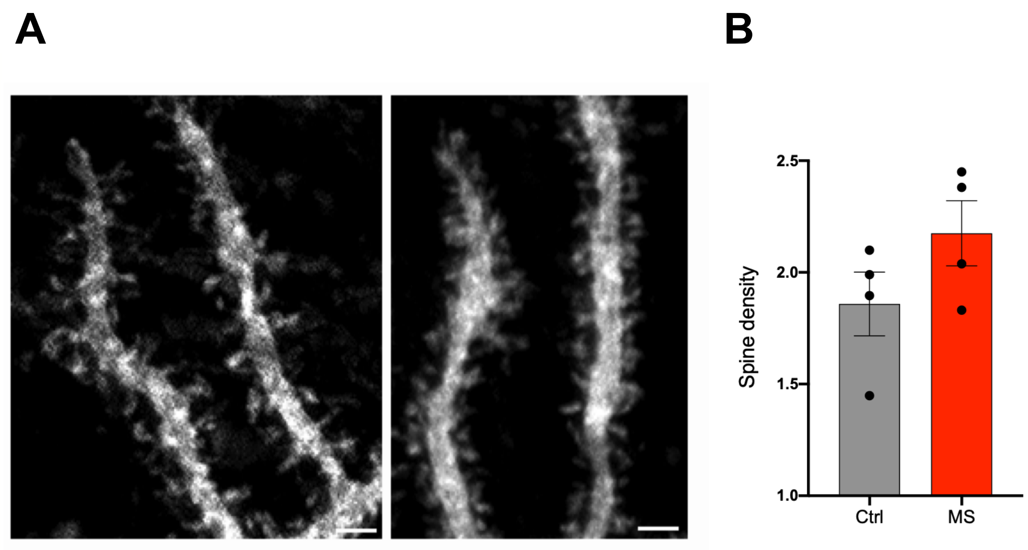


**Figure S1. The number of dendritic spines was unaltered in Purkinje cells (PCs) of maternal separated (MS) mice compared to controls. (A)** Higher magnification images of the dendritic arbor from Purkinje cells of control (left) and MS (right) mice. Scale bar: 10 µm. (**B)** Mean spine density obtained from the analysis of more than 15 dendrites per PC (number of cells: n = 4 control and n = 4 maternal separated PCs).

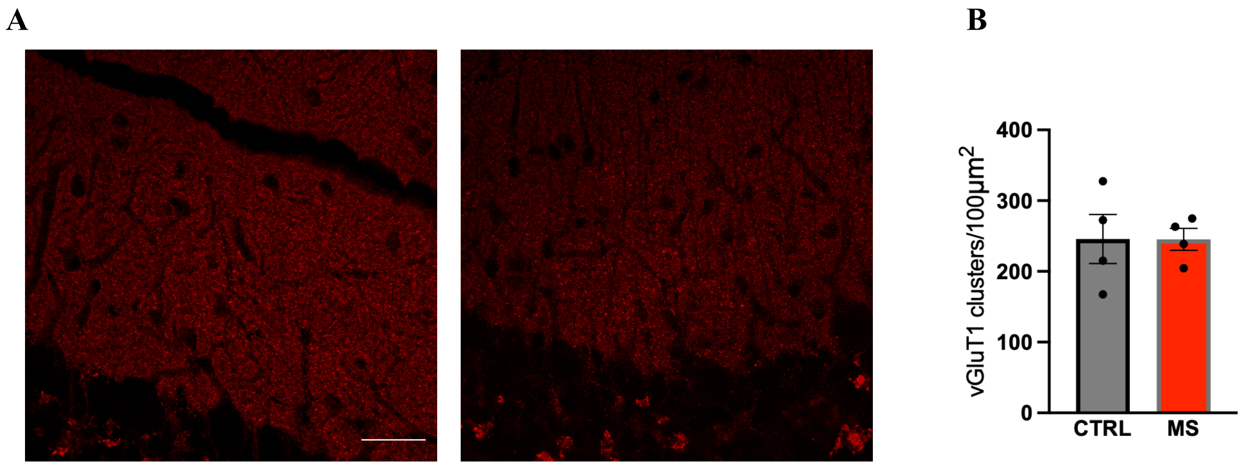


**Figure S2. VGluT1 expression in the molecular layer of the cerebellar cortex. (A)** Representative images showing the expression of VGluT1 in the molecular layer of the cerebellar cortex in control (left) and MS (right) mice. **(B)** Bar graph showing the density of puncta of VGluT1 in both groups of mice. Scale bar: 25 µm. Results are reported as mean ± SEM.

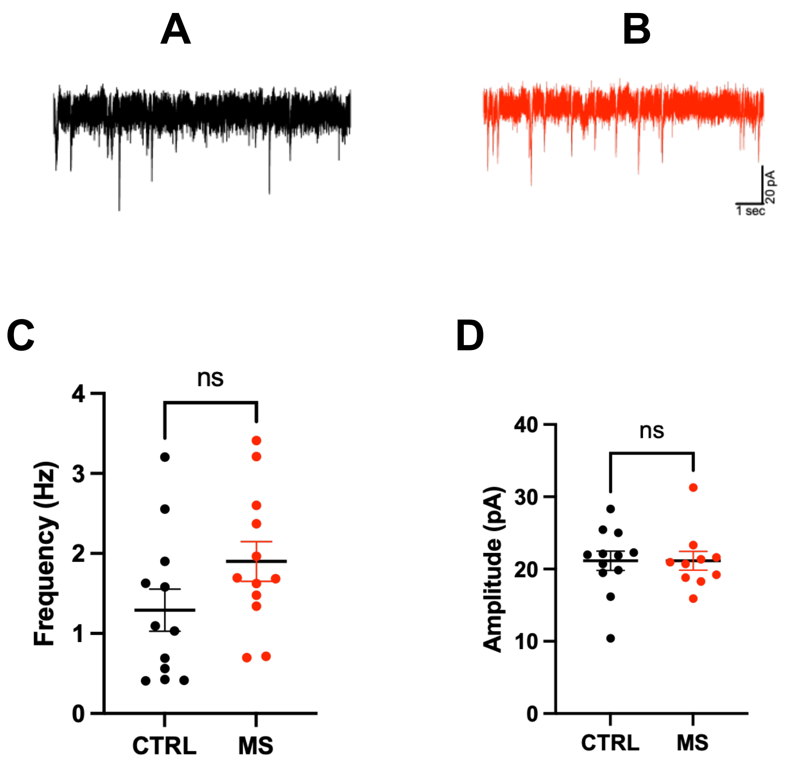


**Figure S3. No significant changes were observed in the frequency or amplitude of spontaneous (s)EPSCs in PCs of MS mice compared to controls.** Representative traces of spontaneous EPSCs from PCs of control (A) and MS (B) mice. Scatter plots showing the frequency (C) and amplitude (D) of sEPSCs in PCs from control and MS mice (n = 11-12 cells for conditions).

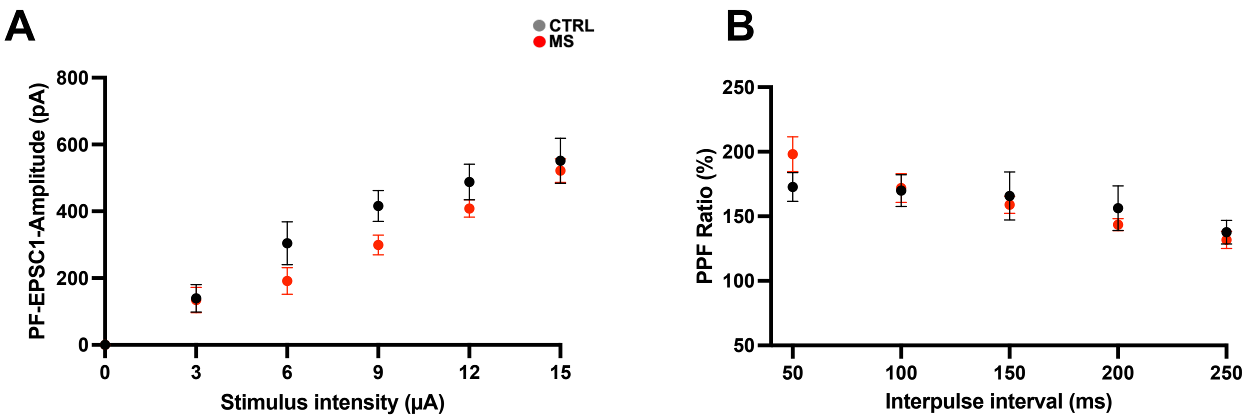


**Figure S4. Input-output relationship and short-term plasticity at parallel fiber-Purkinje cell synapse are not affected in MS mice.** EPSC amplitude elicited by parallel fiber (PF) stimulation increased gradually as the stimulus intensity was increased up to 15μA in both MS and control mice (**A**). The paired-pulse ratio (PPR), which is the ratio of the amplitude of the second response to that of the first, induced by two paired stimuli delivered at inter-pulse intervals ranging from 50 to 250 ms in MS and control mice **(B)**. The magnitude of paired-pulse facilitation at PF-PC synapses in MS mice was not significantly different from that of control mice.

**A**

**B**

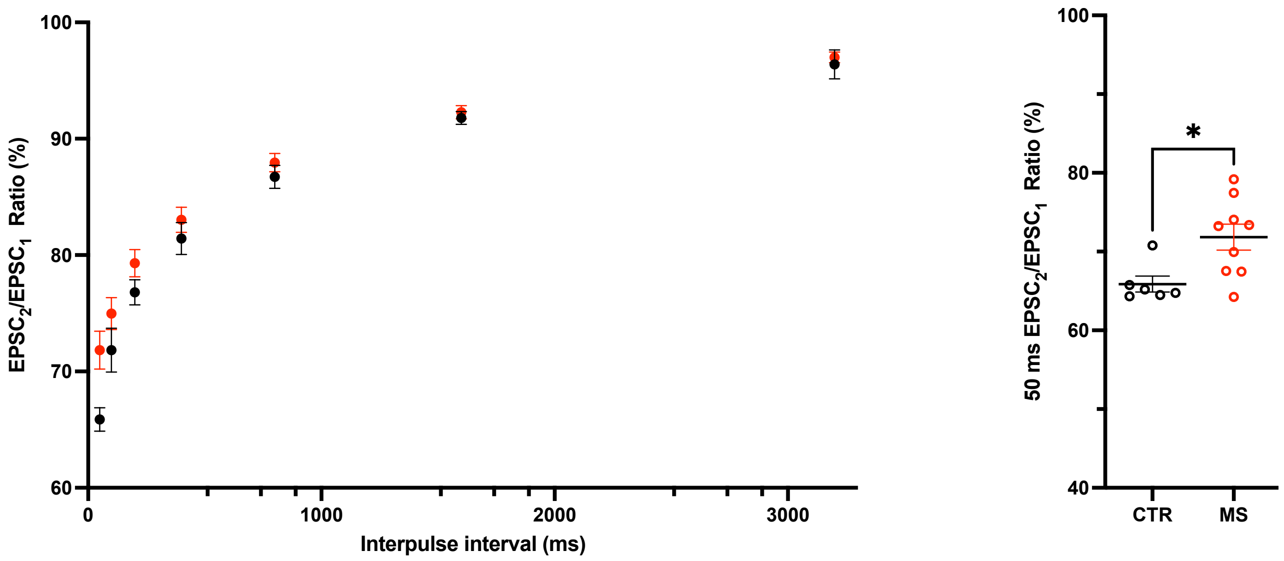


**Figure S5. Short-term plasticity at the climbing fiber-Purkinje cell synapse is affected in MS mice at short interpulse intervals.** Ratio of the amplitude of the second response to that of the first, induced by two paired stimuli of climbing fiber (CF) delivered at inter-pulse intervals ranging from 50 to 3200 ms in MS and control mice **(A)**. The magnitude of paired-pulse depression (PPD) at CF-PC synapses in MS mice was significantly different from that of control mice at the inter-pulse intervals of 50 ms (**B**), unpaired Student's *t*-test. $*p < .05$.

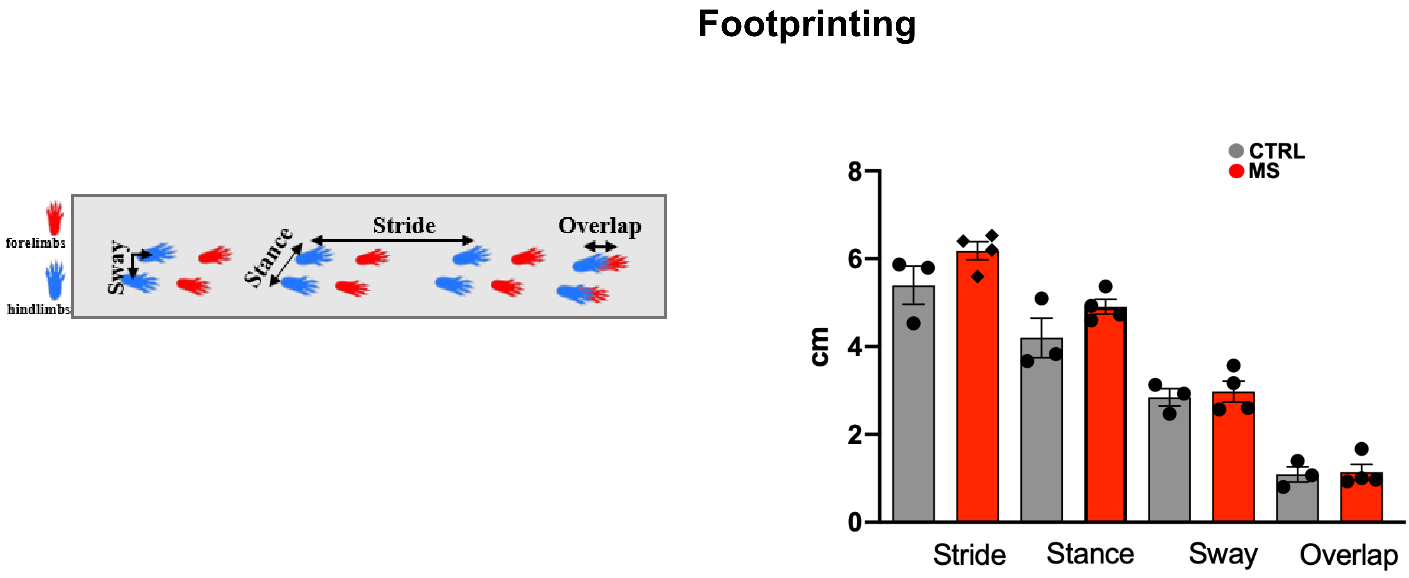


**Figure S6. Footprinting analysis in adult control and MS mice.** On the left, schematic illustration of gait analysis measurements of stride length (distance between subsequent limb placements), stance (distance between left and right limbs), sway (distance between the right and left hindlimb strides) and overlap (between forelimbs and hindlimbs). No significant differences were observed between MS and control animals in any measure of gait analysis, as shown in the bar graph.

**SUPPLEMENTARY MATERIALS**

**Electrophysiological recordings**

Cerebellar slices were prepared as previously described (Lippiello et al, 2020). Animals were anesthetized with isoflurane (Abbott Laboratories, Illinois, USA) and decapitated. The cerebellar vermis was removed and transferred to an ice-cold artificial cerebrospinal fluid (ACSF) containing in mM 125 NaCl, 2.5 KCl, 2 CaCl2, 1 MgCl2, 1.25 NaH2PO4, 26 NaHCO3, and 20 glucose, which was bubbled with 95% $O_2$/5% $CO_2$ (pH 7.4). Parasagittal cerebellar slices (200 μm thick) were obtained using a vibratome (Vibroslice 752, Campden Instruments, Loughborough, UK) and kept for at least 30 min at 32 °C and then at room temperature [67]. Each slice was placed in a recording chamber on an upright microscope (BX50WI, Olympus, Japan) and perfused with oxygenated ACSF at a rate of 2–3 ml/min with ACSF bubbled with t95% $O_2$/5% $CO_2$.

Whole-cell patch-clamp recordings from Purkinje cells (PCs) were performed by an EPC-8 or EPC-10 patch-clamp amplifier (HEKA Elektronik, Lambrecht/Pfalz, Germany). Signals were filtered at 3 kHz, digitized at 10 kz, and stored using the Pulse software (HEKA Elektronik).

For current-clamp recordings, patch pipettes were filled with an internal solution containing in mM: 140 K-gluconate, 10 HEPES, 0.5 EGTA, 4 $MgCl_2$, 4 $Na_2ATP$, 0.4 $Na_3GTP$ and the pH was adjusted to 7.3 with KOH and filtered at 0.2 μm. PCs were held at – 70 mV, and data were filtered at 9.1 kHz and sampled at 20 kHz. A series of depolarizing current steps ranging from 100 to 800 pA in increments of 50 pA, each lasting 500 ms, with step interval of 10 s was delivered to PCs. Gabazine (SR 95531, 20 μM), DAP5 (50 μM) and NBQX (10 μM) were added to the recording chamber to inhibit the $GABA_A$ and ionotropic glutamate receptors of PCs, respectively.

To record the excitatory postsynaptic currents (EPSCs), PCs were held at - 60 mV and parallel fibers were stimulated with an electrode placed into the molecular layer. Double pulses with an interpulse interval of 100 ms were applied every 20 s by a stimulator (A/M Systems, Carlsborg, WA, USA). For long-term potentiation experiments, the patch pipettes were filled with an internal solution containing in mM: 120 potassium gluconate, 9 KCl, 3,5 MgCl2, 10 Hepes, 4 NaCl, 15 sucrose, 4 Na2ATP, 0.4 Na3GTP, 10 EGTA; the pH was adjusted to 7.2 with KOH. For long-term depression experiments, the internal solution contained in mM 130 potassium gluconate, 2 NaCl, 4 MgCl 2, 4Na 2ATP, 0.4 NaGTP, 20 HEPES, 0.25 EGTA. Bicuculline (20 μM) was added to the perfusate to inhibit the GABAA receptors. All data were analyzed using the Igor Pro 6.3 software (Wavemetrics, Lake Oswego, OR, USA). Each EPSC was normalized to the mean EPSC amplitude obtained during 10-min recording before the induction of LTP or LTD.

For construction of input-output curves, the PFs were stimulated at the stimulation intensity ranging from 0 to 15μA. To measure short term plasticity of PF-PC synapse, paired-pulse stimulation of the PF at different intervals (in ms: 50, 100, 150, 200 and 250). Paired-pulse ratio (PPR) was calculated as the ratio of the amplitude of the second (EPSC2) to first response (EPSC1).
We evoked climbing fiber excitatory postsynaptic currents (CF-EPSCs) by delivering square pulses (100 μs) through a stimulating electrode placed in the granular layer. We recorded from PCs at a holding potential of +40 mV. Patch pipettes were filled with an internal solution containing (in mM): 130 CsCl, 4 $MgCl_2$, 10 HEPES, 4 $Na_2ATP$, 0.4 $Na_3GTP$, 10 EGTA, 5 N-(2,6-dimethylphenyl) acetamide-2triethylammonium bromide (QX-314) and the pH was adjusted to 7.3 with CsOH and filtered at 0.2 μm. CF-EPSCs were identified by their all-or-none fashion and the presence of paired pulse depression. When the threshold was detected, twin pulses at different time intervals (50, 100, 150, 200, 400, 800, 1600, 3200 ms) were applied to elicit the paired pulse depression. CF recordings were performed in the presence of the ionotropic $GABA_A$ receptor antagonist, gabazine (SR 95531, 20 μM), in the saline solution.
For spontaneous excitatory postsynaptic currents (sEPSCs), the recording pipettes were filled with an intracellular solution containing, in mM: 130 potassium gluconate, 2 NaCl, 4 $MgCl_2$, 4 $Na_2ATP$, 0.4 $Na_3GTP$, 20 HEPES, and 0.25 EGTA. Bicuculline (10 μM; MedChemExpress) was added to the extracellular solution to block GABAergic inhibitory transmission. Recordings were performed at a holding potential of −60 mV using a MultiClamp 700B amplifier. Signals were digitized at 10 kHz with a Digidata 1440A converter and low-pass filtered at 3 kHz using a Bessel filter controlled by pClamp 11 software (Molecular Devices, Sunnyvale, CA). Continuous 10-min recordings were acquired in gap-free mode, and sEPSC events were detected and analyzed offline using NeuroExpress software v25.7.27.

**Morphology of Purkinje cell dendrites**

Purkinje cells were filled with biocytin through whole-cell patch-clamp pipette and their dendritic arborization was analyzed by performing Sholl analysis. The K-gluconate-based internal solution contained 0.5% biocytin and in mM: 140 K-gluconate, 10 HEPES, 0.5 EGTA, 4 MgCl2, 4 Na2ATP, 0.4 Na3GTP; the pH was adjusted to 7.3 with KOH and the solution was filtered at 0.2 μm. The biocytin-filled PC was visualized with Texas red-conjugated Avidin (1: 1000; Vector, Burlingame, CA, USA). The images were acquired using an E-800 Nikon microscope (Nikon, Melville, NY) connected to a color CCD camera of confocal microscope. The PC dendritic arbor was manually reconstructed and the Sholl analysis was performed using Fiji software. We started our analysis 10 μm from the PC soma with a 22 μm radius. The number of intersections per radius was plotted. For the dendritic spine analysis, we acquired confocal images with a Leica TCS SP5 confocal microscope with a 63x objective (Leica Microsystems, Wetzlar, Germany). The z-stack was acquired at 0.6 μm steps. The analysis was performed with Fiji software, and the spines were counted in the optimal focal planes of the *z*-stack of confocal images. Spine analysis was performed in the distal dendritic arbor of PCs. For each PC, three fields of the dendritic tree were analyzed. The number of spines/μm of dendritic length was provided as spine density.

**Immunostaining**

Sagittal cerebellar slices were incubated in blocking buffer (0.5% Triton X-100, 0.2 % bovine serum albumin, 10% goat serum in PBS) for 4 hours at room temperature. Slices were first incubated with primary antibodies: anti-calbindin (mouse; 1:500 dilution), anti-vGLUT1 (rabbit; 1:500 dilution) or anti-vGLUT2 (guinea pig; 1:1000) at 4°C overnight, followed by fluorescent-conjugated secondary antibodies for 2 hours at room temperature. Images were obtained using a Zeiss LSM 900 confocal microscope at 40x magnification (Carl Zeiss Microscopy GmbH, Germany). Images were acquired at a resolution of 1024 × 1024 pixels, with a scan speed of 8 and a pinhole configuration of 1 Airy unit for each channel. Stacks of images were acquired with a 0.5 μm Z step and reconstructed with Fiji using the maximum intensity projection method (MIP) in the Z-stack function.

Image analysis was conducted using Fiji (ImageJ, NIH). Images were first converted to grayscale, and background noise was reduced using the "Subtract Background" function. Both cerebellar lobes III (anterior) and VIII (posterior) were analyzed. The region of interest (ROI) was defined. The density of vGLUT1 and VGLUT2 within the ROI was quantified using the "Analyze Particles" tool, applying appropriate threshold settings for automated particle counting. Additionally, a subset of images was randomly selected and re-analyzed using the "Cell-counter" tool to validate the automated results (https://doi.org/10.1016/j.celrep.2018.12.069)

**RNA extraction, reverse transcription, and qPCR**

Total RNA was extracted from the cerebellum of P20 control and MS mice using the Trizol reagent, following the manufacturer's protocol. RNA yield and integrity were assessed via spectrophotometric analysis and agarose gel electrophoresis, respectively. For cDNA synthesis, one microgram of RNA was reverse transcribed using the LunaScript® RT SuperMix Kit. Quantitative PCR (qPCR) was conducted with the Luna® Universal qPCR Master Mix on a Bio-Rad Real-Time CFX96 instrument. Each reaction contained cDNA corresponding to 25 ng of total RNA and 0.5 μM specific oligonucleotides listed in table S1. Gene expression data were normalized to HMBS, and expression levels were quantified using the $2^{(-\Delta CT)}$ method, where $\Delta CT$ = CT (gene of interest) - CT (HMBS). All qPCR analyses were performed on at least three biological replicates, with each sample processed independently.

**Table S1. Primers (5'-3') used for qRT-PCR.**

| GENE | Primers Sequence |
|---|---|
| **BDNF** | F: AGCCTCCTCTACTCTTTCTGCT<br>R: GAGTGGCGCCGAACCCTC |
| **CREB** | F: CCCCATTTACCAAACTAGCAG<br>R: CATCCGTACCATTGTTAGCC |
| **GR** | F: CCCCAGGTAAAGAGACAAACGA<br>R: CCTCATTCACGTCCCCTTC |
| **HMBS** | F: ACTGGTGGAGTCTGGAGTCTAGATGGC<br>R: GCCAGGCTGATGCCCAGGTT |

***Footprinting***

To obtain footprints, the front and back pads were coated in red and blue colors, respectively. The animals were then allowed to walk across a white sheet placed on the floor of a runway (4 cm wide and 40 cm long). The assay was performed in a single trial. The gait parameters were measured for 3 consecutive strides. The walking speed and difference in body weight were balanced to exclude differences, which might affect gait parameters. To characterize the walking pattern of each mouse, we measured the distance between subsequent limb placements (stride length), the overlap between forelimbs and hindlimbs, the distance between the left and right limbs (stance width), and the distance between the right and left hindlimb strides (sway distance).
